# Quality of life and satisfaction of users of total tissue-supported and implant-supported prostheses in the municipality of Macapá, Brazil

**DOI:** 10.1101/520197

**Authors:** Éber Coelho Paraguassu, Anneli Celis Mercedes de Cardenas

## Abstract

Despite the high number of total edentulism cases in Brazil, no studies have yet examined the characteristics of people with edentulism in Amapá. Therefore, the objective of this study was to evaluate and compared the satisfaction and quality of life of edentulous users of total tissue-supported and total implant-supported prostheses in Macapá, Amapá, Brazil. Two hundred ninety-nine users of total tissue-supported prostheses and 48 users of total implant-supported prostheses were surveyed using two questionnaires: The Oral Health Impact Profile-14Br and a visual analog scale of satisfaction. The means and standard deviations were used to characterize the quantitative variables and absolute and relative frequencies were used to characterize the qualitative variables as well as certain quantitative variables. When evaluating users’ satisfaction according to the type of prosthesis, we found that users of implant-supported prostheses were 100% satisfied with both upper and lower prostheses. By contrast, among users of tissue-supported prostheses, 90% reported being satisfied with the upper prosthesis, while 56% demonstrated some dissatisfaction with the lower removable prosthesis. All users of implant-supported prostheses reported good quality of life; by contrast, only 5% of users of tissue-supported prostheses reported good quality of life, while 73% reported a reasonable quality of life and 22% a poor quality of life. To our knowledge, this is the first study on this topic in Amapá. The study results are clinically relevant for accurately determining the quality of life of these prosthesis users.

## INTRODUCTION

The number of people with edentulism is growing in Brazil. Edentulism is a physical deficiency related to numerous health problems, such as maxillo-mandibular bone resorption, nutritional deficiencies due to the inability to masticate solid foods, psychological problems, and interpersonal relationship issues.^1^

According to the last National Health Survey—conducted in 2013 by the Brazilian Institute of Geography and Statistics (IBGE) in partnership with the Ministry of Health^2^—roughly 11% of Brazil’s population, or roughly 16 million people, experience complete tooth loss. This proportion was substantially higher among individuals over 60 years, with around 41.5% having lost all their teeth. Another study found that 23% of the Brazilian population had edentulism in one of the archways, while 33% use some form of dental prosthesis.^3^ Given the high prevalence of edentulism, it has become exceedingly important to explore the edentate population’s oral health in order to gain a deeper understanding of their quality of life and how it differs with the specific dental prostheses used.

Research on quality of life and edentulism in Brazil are well advanced; however, they are mostly focused in the south central region of the country. Studies conducted in the Amazon region and more specifically the state of Amapá, which is a state located in the northern end of the country, are limited. The challenges of integration, logistics, communication, and encouragement of the village leave Amapá in a state of darkness, having low scientific production, mainly regarding the quality of life of its population. Therefore, this study aimed to describe the sociodemographic characteristics, satisfaction, and quality of life of edentulous users of total tissue-supported and implant-supported prostheses in the municipality of Macapá, Amapá, Brazil. We hypothesized that the users of implanted prostheses would show better quality of life and greater satisfaction than would users of tissue-supported prostheses. Indeed, we found that the fixation, retention, and stability of the implant-supported prostheses were associated with higher satisfaction and a better quality of life than were the tissue-supported prostheses.

## MATERIALS AND METHODS

### Study population and data collection

A total of 1,393 patients of private clinics and the Centro de Especialidades Odontológicas do Governo do Estado do Amapá (Center for Dentistry Specialties of the Amapá State Government) were screened. Among them, 1,330 were found to be users of the total maxillo-mandibular tissue-supported prosthesis, whereas 63 were users of the total maxillo-mandibular implant-supported prosthesis.

We applied the following inclusion criteria: users with complete edentulism who had up to 10 years of use of either total tissue-supported or total implant-supported prostheses and who lived in Macapá. We excluded the following patients: patients who had used their prostheses for more than 10 years; patients with partial edentulism; patients with complete edentulism only in a single arcade; users of removable partial tissue-supported prosthesis; users of removable or partial dentures on implants; users with complete edentulism who use total tissue-supported prosthesis in one arcade but total implant-supported prosthesis in another; patients who do not live in the urban area of Macapá; and patients with some form of mental or physical disability.

To calculate the ideal sample size for each group, we applied the following formula:

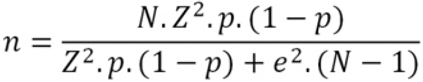

Where n = calculated sample, N = population, Z = standardized normal variable associated with the confidence level, p = true probability of the event, and e = sample error.

We determined the margin of error to be 5% and the confidence level to be 95%. Accordingly, the necessary sample sizes for the total tissue-supported and implant-supported prosthesis users were 299 and 48, respectively.

All potential participants were initially contacted by telephone. Those interested in participating in the study were then visited by the researchers and asked to complete three questionnaires, two of which were validated in past medical literature^4,5^ and were authorized for use by the research ethics committee of the Federal University of Amapá. The study design was approved by this same ethics committee (approval number 2.451.731).

The first questionnaire assessed satisfaction with the prostheses using a visual analog scale (VAS). The questionnaire assessed four aspects—satisfaction with retention, mastication, esthetics, and overall satisfaction—that applied to each of the prostheses that patients used. The VAS for each question ranged from 0 to 10 points (for a potential total score of 40 points). We divided the total score by four to arrive at the final score for satisfaction with the prosthesis. We further categorized the results as follows: 0–2.5 points corresponded to “very dissatisfied,” 2.75–5.0 to “dissatisfied,” 5.25–7.5 to “satisfied,” and 7.75–10 to “very satisfied.”

The second questionnaire was the Oral Health Impact Profile-14 (OHIP-14Br).^6^ This scale contains fourteen questions divided into the following seven subscales:

1. Functional limitations (questions 1 and 2);
2. Physical pain (questions 3 and 4);
3. Psychological discomfort (questions 5 and 6);
4. Physical incapacity (questions 7 and 8);
5. Psychological incapacity (questions 9 and 10);
6. Social incapacity (questions 11 and 12); and
7. Disability (questions 13 and 14).

To calculate patients’ quality of life, we employed the standard calculation method for the OHIP-14Br. First, the following points were assigned to each answer:

- never = 0
- rarely = 1
- sometimes = 2
- frequently = 3
- always = 4

Second, this value was multiplied by a specific weight assigned to each question, as shown in Table 1.

**Table 1:** Weight assigned to each question.

| Question | 1 | 2 | 3 | 4 | 5 | 6 | 7 | 8 | 9 | 10 | 11 | 12 | 13 | 14 |
| --- | --- | --- | --- | --- | --- | --- | --- | --- | --- | --- | --- | --- | --- | --- |
| Weight | 0.51 | 0.49 | 0.34 | 0.66 | 0.45 | 0.55 | 0.52 | 0.48 | 0.6 | 0.4 | 0.62 | 0.38 | 0.59 | 0.41 |

Accordingly, when summing the final scores for the questions, the resultant total scores can range from 0 to 28. Lower scores indicate a better quality of life. More specifically, scores of 0–9.33 indicate a good quality of life, scores of 9.34–18.66 indicate a reasonable quality of life, and scores of 18.67–28 indicate a poor quality of life.

### Statistical analysis

To examine the data, we entered the data in to a spreadsheet in Microsoft Excel 2010, and then transferred the data into SPSS Statistics 22.0 for Windows (IBM. Corp., Armonk, NY) for statistical assessment. We first applied descriptive statistics. The mean and standard deviation were used to describe the continuous variables, and the absolute and relative frequencies were used to describe the categorical as well as certain continuous variables.

For subsequent analyses, we evaluated the normality of the data through the nonparametric Kolmogorov–Smirnov (K-S) test. All the sociodemographic and clinical variables for users of total implant-supported prostheses showed a skewed distribution (*p* > 0.05). By contrast, all the variables for users of total tissue-supported prostheses were normally distributed (*p* < 0.05). Thus, all statistical analyses were conducted using nonparametric tests.

The nonparametric Mann-Whitney U-Test was adopted because it is useful not only for analyzing non-normal data but also for analyzing categorical data. We employed it to compare the mean scores for quality of life, period of use, and satisfaction with the prosthesis between the two prosthesis groups. Spearman’s correlation coefficient was utilized to examine the correlations between quality of life, age, and period of use, as well as quality of life and satisfaction. A 5% significance level was considered for statistical significance. The Spearman’s rho values ranged from null (|ρ = 0|) to perfect (|ρ = 1|).

## RESULTS

Of the 347 participants, 299 used total tissue-supported prostheses and 48 used total implant-supported prostheses. As for the period of use of implant-supported prostheses, the most prevalent answer was one to three years, at 48% (n = 23) for the upper prosthesis and 44% (n = 21) for the lower prosthesis. By contrast, among users of tissue-supported prostheses, four to six years of use was the most prevalent, at 62% (n =185) for the upper prosthesis and 47% (n = 140) for the lower (Table 2).

**Table 2.** Period of Use for the Prostheses.

|  | Implant-supported prosthesis |  |  |  | Tissue-supported prosthesis |  |  |  |
| --- | --- | --- | --- | --- | --- | --- | --- | --- |
| Period of use (years) | Upper prosthesis |  | Lower prosthesis |  | Upper prosthesis |  | Lower prosthesis |  |
|  | n | % | N | % | n | % | N | % |
| 1–3 | 23 | 48 | 21 | 44 | 74 | 25 | 99 | 33 |
| 4–6 | 22 | 46 | 20 | 42 | 185 | 62 | 140 | 47 |
| 7–9 | 3 | 6 | 7 | 14 | 40 | 13 | 60 | 20 |
| ≤10 | 0 | 0 | 0 | 0 | 0 | 0 | 0 | 0 |

\* Data analyzed in SPSS Statistics 22.0 for Windows (IBM. Corp., Armonk, NY)

When comparing the prostheses according to period of use, with regard to the upper prosthesis, the tissue-supported prostheses had a greater period of use (mean = 4.63 ±1.64 years) than did the implant-supported prostheses (*p* < 0.001). As for the lower prosthesis, the difference in the period of use between the two types of prostheses was minor and no significant (*p* > 0.05) (Table 3).

**Table 3.**
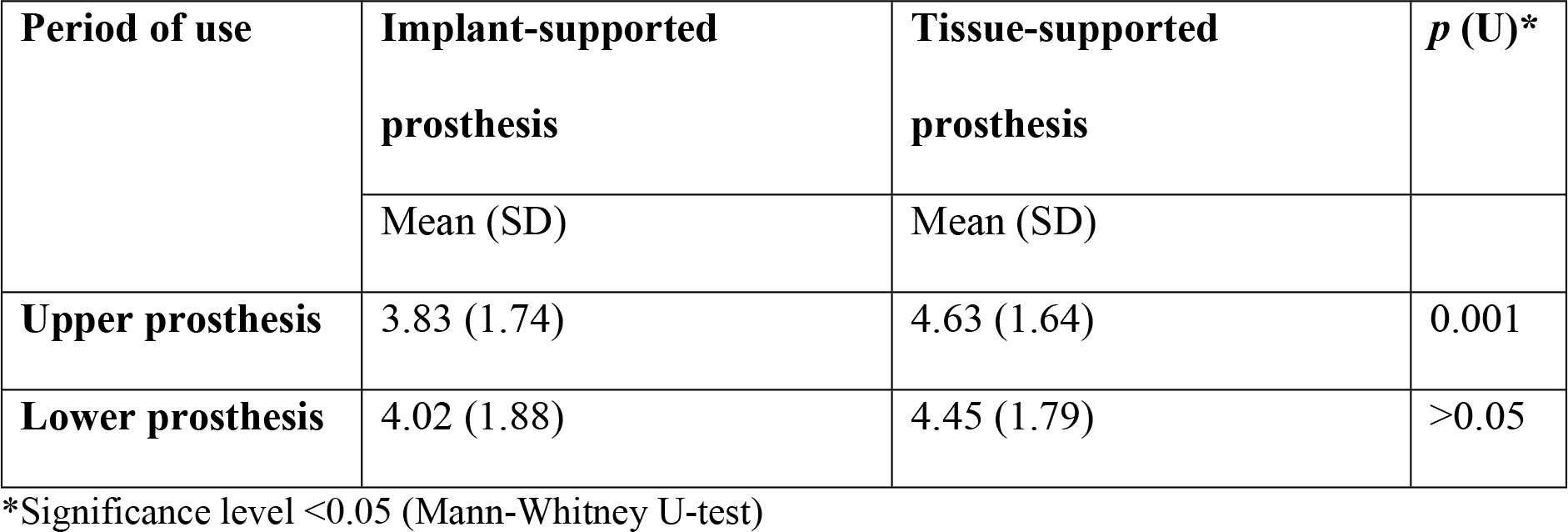
Means and Standard Deviations of the Period of Use of the Prostheses.

| Period of use | Implant-supported prosthesis | Tissue-supported prosthesis | $p$ (U)* |
| --- | --- | --- | --- |
|  | Mean (SD) | Mean (SD) |  |
| Upper prosthesis | 3.83 (1.74) | 4.63 (1.64) | 0.001 |
| Lower prosthesis | 4.02 (1.88) | 4.45 (1.79) | >0.05 |

\*Significance level <0.05 (Mann-Whitney U-test)

All patients with implant-supported prostheses reported being very satisfied with their lower and upper prostheses (100%; n = 48). As for tissue-supported prostheses, 90% (n = 269) of the patients reported feeling very satisfied with the upper prosthesis, while 56% displayed some dissatisfaction with the lower prosthesis (Table 4).

**Table 4.** Patients’ Satisfaction with the Prostheses (Visual Analogue Scale)

| Classification | Implant-supported prosthesis |  |  |  | Tissue-supported prosthesis |  |  |  |
| --- | --- | --- | --- | --- | --- | --- | --- | --- |
|  | Upper prosthesis |  | Lower prosthesis |  | Upper prosthesis |  | Lower prosthesis |  |
|  | N | % | N | % | N | % | N | % |
| **Very satisfied** | 48 | 100 | 48 | 100 | 28 | 9 | 3 | 1 |
| **Satisfied** | 0 | 0 | 0 | 0 | 269 | 90 | 112 | 38 |
| **Dissatisfied** | 0 | 0 | 0 | 0 | 2 | 1 | 168 | 56 |
| **Very dissatisfied** | 0 | 0 | 0 | 0 | 0 | 0 | 16 | 5 |

\* Data analyzed in SPSS Statistics 22.0 for Windows (IBM. Corp., Armonk, NY)

The mean VAS satisfaction score was significantly greater (*p* < 0.01) for implant-supported prostheses, with a mean of 9.39 (±0.50) for the upper and 9.47 (±0.48) for the lower (Table 5).

**Table 5.** Means and Standard Deviations of Patients’ Satisfaction with the Prostheses.

| VAS Satisfaction | Implant-supported prosthesis | | Tissue-supported prosthesis | | $p (U)^*$ |
| --- | --- | --- | --- | --- | --- |
|  | Mean | SD | Mean | SD |  |
| **Upper prosthesis** | 9.39 | 0.50 | 6.63 | 0.71 | 0.01 |
| **Lower prosthesis** | 9.47 | 0.48 | 4.82 | 1.11 |  |

\*Significance level $< 0.05$ (Mann-Whitney U-test)

All users of implant-supported prostheses reported having good quality of life (100%; n = 48). As for the users of tissue-supported prostheses, 73% (n = 220) reported reasonable quality of life; 22% (n = 65) reported a poor quality of life; and only 5% (n = 14) reported a good quality of life (Table 6).

**Table 6.** Quality of Life Classification (OHIP-14Br Questionnaire) for Patients Who Used Prostheses.

| Classification of quality of life | Implant-supported prosthesis |  | Tissue-supported prosthesis |  |
| --- | --- | --- | --- | --- |
|  | N | % | N | % |
| **Good** | 48 | 100 | 14 | 5 |
| **Reasonable** | 0 | 0 | 220 | 73 |
| **Poor** | 0 | 0 | 65 | 22 |

\* Data analyzed in SPSS Statistics 22.0 for Windows (IBM. Corp., Armonk, NY)

The mean quality of life score of patients who used implant-supported prostheses was 3.24 (±01.32), which was significantly lower (*p* < 0.0001) than that of patients who used tissue-supported prostheses, who had a mean score of 15.63 (±3.23). Thus, individuals who used the implant-supported prostheses had a better quality of life than did those who used tissue-supported prostheses (Table 7).

**Table 7.** Means and Standard Deviations of Quality of Life of Patients Who Used Prostheses.

| Quality of life evaluation | Mean | SD | *p (U)* |
| --- | --- | --- | --- |
| **Implant-supported prosthesis** | 3.24 | 1.32 | <0.0001 |
| **Tissue-supported prosthesis** | 15.63 | 3.23 |  |

\*Significance level <0.05 (Mann-Whitney U-test)

We found strong correlations of quality of life with age and period of use for tissue-supported prosthesis users. Specifically, the longer the use of the prosthesis and the greater their age, the worse was their quality of life. These relationships did not exist among users of implant-supported prostheses (Table 8). No other correlation showed statistical significance (*p* > 0.05).

**Table 8.** Correlations between Quality of Life, Age, and Period of Use.

|  | Age | Period of use |  |
| --- | --- | --- | --- |
|  |  | Longer | Shorter |
| Quality of life for implant-supported prosthesis | 0.2 ($p > 0.5$) | 0.2 ($p > 0.05$) | 0 |
| Quality of life for tissue-supported prosthesis | 0 | 0.7 ($p < 0.0001$) | 0.9 ($p < 0.0001$) |

\* Data analyzed in SPSS Statistics 22.0 for Windows (IBM. Corp., Armonk, NY)

Among users of implant-supported prostheses, quality of life was negatively correlated with satisfaction with the upper prosthesis (ρ *=* −0.3, *p* < 0.05). Thus, better quality of life was associated with greater satisfaction with the prosthesis. A similar trend was found for tissue-supported prostheses, for both the upper (ρ = −0.5, *p* < 0.0001) and lower (ρ = −0.7, *p* < 0.0001) protheses. This relationship was not statistically significant for lower implant-supported prostheses (Table 9). No other correlation was significant (*p* > 0.05).

**Table 9.** Correlation between Quality of Life and Satisfaction (Visual Analogue Scale)

|  | Implant-supported satisfaction |  | Tissue-supported satisfaction |  |
| --- | --- | --- | --- | --- |
|  | Upper prosthesis | Lower prosthesis | Upper prosthesis | Lower prosthesis |

| | $\rho$ ($p$) | $\rho$ ($p$) | $\rho$ ($p$) | $\rho$ ($p$) |
| --- | --- | --- | --- | --- |
| **Quality of life of implant** | -0.3<br>($<0.05$) | -0.1<br>($>0.05$) | - | - |
| **Quality of life of removable prosthesis** | - | - | -0.5<br>($<0.0001$) | -0.7<br>($<0.0001$) |

\* Data analyzed in SPSS Statistics 22.0 for Windows (IBM. Corp., Armonk, NY)

## DISCUSSION

This study has implications for the quality of life of the people of the Brazilian Amazon region, mainly the state of Amapá (which is representative of this region), as it is the only study to our knowledge to examine the quality of life and satisfaction of individuals with complete edentulism who are using total dental prostheses in this region. The lack of research in this region is a cause for concern, as the Amazon region occupies about 40% of Brazilian territory (3.5 million km²) and has a population of approximately 25 million people according to the latest survey by the IBGE.^7^ This study can therefore provide a basis for future studies on the quality of life of the inhabitants of this region and thereby contribute to the international literature.

When looking at the period of use of implant-supported prostheses, similar results were observed between the 1–3 years group and the 4–6 years group, both for the upper prosthesis (48% and 46%, respectively) and lower prosthesis (44% and 42%). Only 6% and 14% of patients had used the upper and lower prostheses, respectively, for 7 to 9 years. The mean period of use of upper implant-supported prostheses was 3.83 years, while the mean period for lower prostheses was 4.02 years. The users of upper and lower total tissue-supported prostheses were more prevalent in the 4–6 years group, corresponding to 62% and 47% of the total users and having a mean of 4.63 years and 4.45 years, respectively. Notably, the period of use of upper and lower prostheses did not differ significantly by type of prosthesis. The literature shows contradictory results on this, with some studies corroborating the results of the present study and others not doing so.^8–13^ This leads us to hypothesize that there is no regularity in the period of use of the prostheses according to country or region.

The level of satisfaction with the prostheses differed between the two groups. All the users of implant-supported prostheses were very satisfied with their prosthesis, with a mean score of about 9.39 for the upper prosthesis and 9.47 for the lower prosthesis. The users of tissue-supported prosthesis were less satisfied, with a mean score of 6.63 for the upper prosthesis (and thus classified as satisfied) and 4.82 for the lower prosthesis (classified as dissatisfied). This difference can be attributed to the fact that implant-supported prostheses are fixed and have greater retention and stability, thereby providing the user with greater masticatory efficiency and confidence, and enabling the reestablishment of interpersonal relations; these in turn can lead to greater well-being and improved satisfaction.^14–16^

Quality of life also significantly differed between the two groups. All users of implant-supported prosthesis had good quality of life (with a mean score of 3.24 on the OHIP-14Br). Patients with tissue-supported prosthesis had a mean score of 15.33, with most having a reasonable quality of life (73%) and 22% showing poor quality of life; only 5% showed good quality of life. We also found that quality of life is directly associated with period of use among users of tissue-supported prostheses, as well as with satisfaction. Users in general who were more satisfied with their prostheses also tended to have a better quality of life. In addition, users of tissue-supported prostheses showed a significantly decreasing quality of life as the length of use of their prosthesis increased, especially for the lower prosthesis. The fixation and stability of implant-supported prostheses, in addition to improving satisfaction, increases the quality of life of their users because they restore aesthetic, phonetic, and masticatory functions almost completely, thus restoring people’s social life and reducing or eliminating any concerns with their teeth.^17^

Consideration of bone resorption in people with complete edentulism is extremely important for understanding the quality of life of users of total tissue-supported prostheses. A high rate of maxillo-mandibular bone resorption is typically observed in the first year after extractions in people with total teeth loss. This resorption can reach 12 mm in that first year, after which it tends to stabilize to 1– 2 mm/year until complete resorption of the alveolar ridge is achieved. The continuous process of bone resorption causes the bearing area to become very thin and small, eventually to the point where the tissue-supported prostheses cannot fit and lose all or a large part of their retention. Thus, this makes it difficult for the patient to feed, speak, or express any facial gestures without displacing the prosthesis.^18^ Continuous bone resorption may be the reason for the association between the longer use of tissue-supported prosthesis and the lower satisfaction and quality of life.

Resorption takes place differently in the mandible and maxilla. In a comparative study of bone resorption between these areas,^19^ it was observed that the mandible tends to reabsorb bone about 25% faster than the maxilla. The mandible is a highly critical region for total tissue-supported prostheses, because, besides this faster resorption rate, it has a bearing area 1.8% lower than the maxilla; this lowers the stability and retention of the prosthesis. In most cases, bone resorption reaches a point where the prosthesis becomes effectively useless after few years.^20^ This might also explain why the satisfaction with lower tissue-supported prostheses was lower than was that of the upper prostheses (which in turn led to lower quality of life).

This study has some limitations. First, we did not collect data from other states of the Amazon region, due to the difficulties in interstate logistics imposed by the forest—specifically, there is no terrestrial integration between the states and we observed a lack of collaboration for participation in the study by populations in other states.

Nevertheless, our findings can guide public policies so that Brazil’s unique health system can offer its users the possibility of fixation of removable prostheses through dental implants, rather than relying on removable prostheses, as is done today.

## Conclusions

Users of implant-supported prostheses had a significantly better quality of life, and greater satisfaction with their prostheses than did users of total tissue-supported prostheses. We found direct relationships between the period of use of total tissue-supported prostheses and quality of life and satisfaction. Specifically, the longer the period of use, the lower the satisfaction and quality of life. Bone resorption may be a major contributor to the lower quality of life and satisfaction among users of total tissue-supported prosthesis. Fixation of total prostheses using implants brings greater stability and retention, and consequently better masticatory function, to its users, which seems to lead to a considerable increase in users’ quality of life and satisfaction with the prosthesis.

## Acknowledgements

The authors wish to thank the entire graduate program of health sciences at the Federal University of Amapá, especially our counselor Dr. Anneli Celis de Cardenas. We are also thankful for the coordination of oral health department of the state of Amapá, especially Dr. Rodrigo Vançan, who authorized the data collection of almost all patients with tissue-supported prosthesis, and OdontoImplantes Dental Clinic, which authorized the data collection of almost all the patients with implant-supported prosthetics.

## References

1. Goiato MC, Torcato LB, dos Santos DM, Moreno A, Antenucci RM, de Carvalho Dekon SF. Quality of life and satisfaction of patients wearing implant-supported fixed partial denture: A cross-sectional survey of patients from Araçatuba city, Brazil. Clin Oral Implants Res. 2015;26(6):701–708. Accessed September 27, 2018.

2. Brazilian Institute of Geography and Statistics (IBGE) in partnership with the Ministry of Health in 2013. National Health Survey (PNS). Available at: ftp://ftp.ibge.gov.br/PNS/2013/pns2013.pdf. Accessed 09, 27, 2018.

3. Nico LS. Saúde bucal autorreferida da população adulta brasileira: resultados da Pesquisa Nacional de Saúde 2013. Ciênc Saúde Colet. 2016;21(2). Accessed September 30, 2018.

4. Silva ME, Villaça ÊL, Magalhães CS, Ferreira EF. Impacto da perda dentária na qualidade de vida. Ciênc Saúde Colet. 2010;15:841–850. Accessed September 30, 2018.

5. Slade GD, Spencer AJ. Development and evaluation of the Oral Health Impact Profile. Community Dent Health. 1994;11(1):3–11. Accessed September 30, 2018.

6. Oliveira BH, Nadonovsky P. Psychometric properties of the Brazilian version of the Oral Health Impact Profile-Short Form. Community Dent Oral Epidemiol. 2005;33(4):307–314. Accessed October 02, 2018.

7. Insituto Brasileiro de Geografia e Estatistica (IBGE). Censo demográfico 2010. Available at: https://www.ibge.gov.br/estatisticas-novoportal/sociais/populacao/9662-censo-demografico-2010.html?=&t=o-que-e. Accessed October 02, 2018.

8. Silva, Maria Elisa de Souza, et al. “Impacto da perda dentária na qualidade de vida.” Ciência & Saúde Coletiva 15 (2010): 841–850. Accessed October 15, 2018.

9. Azevedo M, Correa MB, Azevedo JS, Demarco FF. Dental prosthesis use and/or need impacting the oral health-related quality of life in Brazilian adults and elders: Results from a National Survey. J Dent. 2015;43(12):1436–1441. Accessed October 19, 2018.

10. Ilha L, Martins AB, Abegg C. Oral impact on daily performance: need and use of dental prostheses among Brazilian adults. J Oral Rehab. 2016;43(2):119–126. Accessed October 19, 2018.

11. Machado FW, Perroni AP, Nascimento GG, Goettems ML, Boscato N. Does the sense of coherence modifies the relationship of oral clinical conditions and oral health-related quality of life? Qual Life Res. 2017;26(8):2181–2187. Accessed October 19, 2018.

12. Castrejón?Pérez, Borges?Yáñez SA, Irigoyen-Camacho ME, Cruz-Hervert LP. Negative impact of oral health conditions on oral health related quality of life of community dwelling elders in Mexico city, a population based study. Geriatr Gerontol Int. 2017;17(5):744–752. Accessed October 21, 2018.

13. Goncalves TM, Campos CH, Garcia MR. Effects of implant-based prostheses on mastication, nutritional intake, and oral health-related quality of life in partially edentulous patients: A paired clinical trial. Int J Oral Maxillofac Implants. 2015;30(2): 391–396. Accessed October 21, 2018.

14. Awad MA, Lund JP, Shapiro SH, et al. Oral health status and treatment satisfaction with mandibular implant overdentures and conventional dentures: a randomized clinical trial in a senior population. Int J Prosthodont. 2003;16(4):390–396. Accessed October 22, 2018.

15. Lang LA, Garcia LT, Teich ST, Olvera N. Comparison of the impact on quality of life of immediate versus delayed implant placement supporting immediately loaded mandibular overdenture. Refuat Hapeh Vehashinayim (1993). 2016;33(1):6–14, 59. Accessed October 23, 2018.

16. Preciado A, Del Río J, Lynch CD, Castillo-Oyagüe R. A new, short, specific questionnaire (QoLIP-10) for evaluating the oral health-related quality of life of implant-retained overdenture and hybrid prosthesis wearers. J Dent. 2013;41(9):753–763. Accessed October 23, 2018.

17. Yunus N, Masood M, Saub R, et al. Impact of mandibular implant prostheses on the oral health?related quality of life in partially and completely edentulous patients. Clin Oral Implant Res. 2016;27(7):904–909. Accessed October 23, 2018.

18. Assunção WG, Zardo GG, Delben JA, Barão VA. Comparing the efficacy of mandibular implant-retained overdentures and conventional dentures among elderly edentulous patients: satisfaction and quality of life. Gerodontology. 2007;24:235–238. Accessed 10, 23, 2018

19. Koshino H, Hirai T, Yokoyama Y, et al. Mandibular residual ridge shape and masticatory ability in complete denture wearers. J Jpn Prosthodont. 2008;52:488–493. Accesed October 23, 2018.

20. Nunez CMNO. Efetividade de um protocolo de tratamento simplificado com próteses totais sobre a satisfação dos pacientes com as próteses e qualidade de vida relacionada à saúde buccal [Dissertation]. Catalão: Universidade Federal de Goiás 2011. Accessed October 24, 2018.

